# E-cadherin represses anchorage-independent growth in sarcomas through both signaling and mechanical mechanisms

**DOI:** 10.1101/347815

**Authors:** Mohit Kumar Jolly, Kathryn E. Ware, Shengnan Xu, Shivee Gilja, Samantha Shetler, Yanjun Yang, Xueyang Wang, R. Garland Austin, Daniella Runyambo, Alexander J. Hish, Suzanne Bartholf DeWitt, Jason T. George, R. Timothy Kreulen, Mary-Keara Boss, Alexander L. Lazarides, David L. Kerr, Drew G. Gerber, Dharshan Sivaraj, Andrew J. Armstrong, Mark W. Dewhirst, William C. Eward, Herbert Levine, Jason A. Somarelli

## Abstract

E-cadherin, an epithelial-specific cell-cell adhesion molecule, plays multiple roles in maintaining adherens junctions, regulating migration and invasion, and mediating intracellular signaling. Downregulation of E-cadherin is a hallmark of epithelial-mesenchymal transition (EMT) and correlates with poor prognosis in multiple carcinomas. Conversely, upregulation of E-cadherin is prognostic for improved survival in sarcomas. Yet, despite the prognostic benefit of E-cadherin expression in sarcoma, the mechanistic significance of E-cadherin in sarcomas remains poorly understood. Here, by combining mathematical models with wet-bench experiments, we identify the core regulatory networks mediated by E-cadherin in sarcomas, and decipher their functional consequences. Unlike in carcinomas, E-cadherin overexpression in sarcomas does not induce a mesenchymal-epithelial transition (MET). However, E-cadherin acts to reduce both anchorage-independent growth and spheroid formation of sarcoma cells. Ectopic E-cadherin expression acts to downregulate phosphorylated CREB (p-CREB) and the transcription factor, TBX2, to inhibit anchorage-independent growth. RNAi-mediated knockdown of TBX2 phenocopies the effect of E-cadherin on p-CREB levels and restores sensitivity to anchorage-independent growth in sarcoma cells. Beyond its signaling role, E-cadherin expression in sarcoma cells can also strengthen cell-cell adhesion and restricts spheroid growth through mechanical action. Together, our results demonstrate that E-cadherin inhibits sarcoma aggressiveness by preventing anchorage-independent growth.

## Introduction

Sarcomas – deadly cancers that arise from tissues of a mesenchymal lineage – are highly aggressive, with five year survival rates of just 66% (1). Despite their mesenchymal origin, some sarcomas undergo phenotypic plasticity in which they gain “epithelial-like” traits (2–4). While this transition to a more epithelial-like state is now being recognized as a feature of multiple subtypes of soft tissue sarcoma and osteosarcoma (2–4), there are also a number of sarcoma subtypes that are classically known to exhibit epithelioid features pathologically, including synovial sarcoma (5), epithelioid sarcoma (6), and adamantinoma (7). One might expect the acquisition of epithelial-like traits to be of little relevance in mesenchymal tumors, yet that is not the case. Phenotypic plasticity is clinically important in sarcoma patients: Sarcoma patients whose tumors express epithelial-like biomarkers have improved outcomes relative to patients with more “mesenchymal-like” tumors (2–4,8).

Phenotypic plasticity observed in sarcomas is reminiscent of the phenomenon of epithelial plasticity in carcinomas. Epithelial plasticity refers to reversible transitions between epithelial and mesenchymal phenotypes. In carcinomas, the phenotypic transition to a more mesenchymal-like state via an epithelial-mesenchymal transition (EMT) promotes migratory and invasive gene expression programs that facilitate cancer cell invasion and metastatic seeding (9). Subsequent to metastatic dissemination, a reversion to an epithelial-like state via mesenchymal - epithelial transition (MET) re-awakens proliferative signals within the metastatic niche to enable metastatic colonization (9).

In carcinomas, the gene expression programs that control EMT/MET are regulated at multiple stages, including through epigenetics (10), transcription (11), microRNAs (12), alternative splicing (13,14), and post-translational protein stability (15). These regulatory mechanisms control genes involved in cell polarity, cytoskeletal architecture, cell-substrate adhesion, and cell-cell adhesion. One of these genes, E-cadherin, is an epithelial-specific cell-cell adhesion molecule that has multiple functions in maintenance of adherens junctions (16), cytoskeletal organization (17), migration (18,19), and intracellular signaling (20). Downregulation of E-cadherin is a marker of poor prognosis in multiple cancers of an epithelial origin (21,22). In addition, loss-of-function germline mutations in E-cadherin predispose individuals to familial gastric cancer (23), early onset colorectal cancer (24), and hereditary lobular breast cancer (25).

Consistent with its known tumor suppressor role in carcinomas, E-cadherin upregulation is also prognostic for improved survival in sarcomas (8). However, despite the prognostic importance of E-cadherin in sarcomas, little is known about the molecular mechanisms that underlie improved outcomes of E-cadherin upregulation in mesenchymally-derived cancers. Here, we use a combined theoretical-experimental approach to decipher the gene regulatory networks driven by E-cadherin in sarcomas. Though not a generalized phenomenon, in some carcinomas E-cadherin is sufficient to induce a more epithelial-like phenotype (26); however, our study demonstrates E-cadherin expression is not sufficient to alter epithelial plasticity biomarkers, migration, or invasion. E-cadherin expression did, however, significantly inhibit both anchorage-independent growth and spheroid growth in sarcoma cells. Non-cancer cells that become detached from the normal tissue architecture undergo a cell death program known as anoikis. Resistance to anoikis is a hallmark of cancer progression and of an aggressive phenotype. E-cadherin-mediated repression of anchorage-independent growth was accompanied by downregulation of phospho-CREB and the transcription factor, TBX2. TBX2 knockdown led to reductions in total and phospho-CREB and phenocopied E-cadherin-mediated inhibition of anchorage-independent growth. In addition to its signaling role, E-cadherin repressed spheroid size through increased cell-cell adhesion. Together, these results indicate that E-cadherin acts through both signaling and mechanical roles to inhibit sarcoma aggressiveness by suppressing both anchorage-independent growth and spheroid growth.

## Materials and Methods

### Analysis of the prognostic impact of E-cadherin in clinical data sets

The prognostic significance of E-cadherin in osteosarcoma was analyzed using the R2: Genomics Analysis and Visualization Platform (https://hgserver1.amc.nl/cgi-bin/r2/main.cgi). Kaplan Meier plots of metastasis-free survival and overall survival were constructed using the Kaplan Meier option with the ‘mixed osteosarcoma’ dataset (27). For soft tissue sarcomas, Kaplan Meier plots were constructed in JMP13.0 as previously described (Somarelli et al. 2016) using E-cadherin RNA-Seq and protein levels from The Cancer Genome Atlas (TCGA). Correlations between E-cadherin levels and EMT score were performed using RNA-Seq data from TCGA and analyzed in JMP13.0.

To calculate EMT scores, raw TCGA sarcoma gene expression data for 259 samples was accessed from the R2: Genomics Analysis and Visualization Platform (http://r2.amc.nl). EMT quantification was applied by using a small set of EMT-predictors in addition to a normalization signature of EMT-uncorrelated transcripts as described previously (28). Each sample was assigned an EMT score, 0 < μ < 2, with E scores μ_E_ < 0.5, M scores μ_M_ > 15 and hybrid E/M scores 0.5 ≤ μ_E/M_ ≤ 15 intermediary to both E and M scores. Samples were partitioned into E-cadherin^low^ and E-cadherin^high^ categories based on median E-cadherin expression. The probability distribution for each group was estimated by spline interpolation of the empirical EMT score histogram.

### Cell culture conditions, ectopic E-cadherin expression, and flow cytometry

143B, U2OS, Abrams, and RD cells were cultured in high glucose Dulbecco’s Modified Eagle’s Medium (DMEM) supplemented with 10% fetal bovine serum (FBS) and 1% penicillin/streptomycin (pen-strep) at 37°C and 5% CO_2_ in a humidified incubator. 143B, U2OS, and RD cells were obtained from the Duke University Cell Culture Facility, which performs routine mycoplasma testing and verifies cell identity by analysis of short tandem repeats. Abrams cells were a gift of Dr. Douglas Thamm (Colorado State University). To generate sarcoma cells stably expressing E-cadherin, 143B, U2OS, and RD cells were stably transfected with an empty vector (pcDNA3) or hE-cadherin-pcDNA3 (a gift from Barry Gumbiner; Addgene plasmid # 45769; (29)). E-cadherin positive cells were sorted by flow cytometry using a PerCP710 anti-E-cadherin antibody (eBioscience cat. 46-3249). To do this, confluent cells in a T75 flask were washed with phosphate buffered saline (PBS), scraped, pelleted by centrifugation at 250 x g for 10 minutes, resuspended in 5% bovine serum albumin in PBS, incubated for 30 minutes, centrifuged as above, resuspended in anti-E-cadherin antibody dilution at 5 μL in 100 μL of DMEM with 5% FBS per 1 x 10^6^ cells, incubated for 30 minutes, centrifuged again, washed with 5 ml of PBS, spun, resuspended in 0.5 ml of DMEM supplemented with 5% FBS, and filtered through a 30 μM filter (Partek) into a flow cytometry tube. E-cadherin+ cells were sorted into fully supplemented DMEM at the Duke University Flow Cytometry Shared Resource.

### Knockdown by siRNAs

A total of 20 nM of Allstars non-silencing (Qiagen) or target gene siRNA (Qiagen) was diluted in Opti-MEM and mixed with Lipofectamine RNAiMax diluted in Opti-MEM. The RNAiMax-siRNA mixtures were incubated for 20 minutes at room temperature, followed by plating of the mixtures in 24-well plates. A total of 25,000 cells/well diluted in fully supplemented DMEM were seeded atop the RNAiMax-siRNA mixtures and incubated overnight at 37°C and 5% CO_2_ in a humidified incubator. Media was replaced the next day. After 72 hours of incubation, cells were collected in the appropriate buffer or fixative for downstream applications.

### Gene expression analysis by qRT-PCR

RNA was extracted from cultured cells using the Zymo Quick RNA Miniprep Kit following the manufacturer’s protocol. Reverse transcription (RT) reactions were performed as described (Somarelli et al. 2016) using the ABI High Capacity cDNA Reverse Transcription Kit (ThermoFisher). RT reactions were incubated following the manufacturer’s protocol in a SimpliAmp thermocycler (Life Technologies). RT reactions were diluted 1:5 in nuclease-free H2O, and RT-qPCR was performed as described (Somarelli et al. 2016) in a Vii7 real time-PCR detection system (Applied Biosystems).

### Western blotting, immunofluorescence staining, phospho-kinase arrays, and ELISAs

Western blotting and immunofluorescence staining were performed as previously described (Somarelli et al. 2016). A complete list of primary antibodies and their dilutions is provided in the Supplementary Text. To prepare lysates for phospho-kinase arrays, E-cadherin^−^ and E-cadherin^+^ cells were sorted by flow cytometry prior to each assay, seeded into 10 cm dishes, and allowed to grow for 48-72 hours until they reached 80% confluence. Cells were then washed twice with ice-cold PBS and scraped gently in 2 ml of PBS using cell scrapers. Cells were counted using the CountessII system (Life Technologies), and cells were solubilized at 1 x 10 cells/ml in Lysis Buffer 6 provided as part of the Proteome Profiler Human Phospho-Kinase Array Kit (R&D Systems). Total protein concentrations in 143B+empty vector and 143B+E-cadherin lysates were estimated using a combination of both Bradford assays and western blotting (to analyze consistent loading of intact protein with β-actin as a loading control). Lysates were used in the Proteome Profiler Human Phospho-Kinase Array (R&D cat. ARY003b) according to the manufacturer’s instructions. Membranes were exposed to x-ray films and visualized on a light box. The same procedure was used to generate lysates for ELISAs, and p-CREB levels were quantified using DuoSet IC Human/Mouse/Rat Phospho-CREB (S133) ELISA kits (R&D Systems).

### Migration, invasion, proliferation, soft agar, spheroid growth assays and cell-cell adhesion assays

For migration and invasion assays, a total of 500 μL of DMEM (with 10% FBS and 1% pen/strep) was added to the top and bottom of Boyden chamber wells for two hours to hydrate the transwell membranes. Next, 50,000 cells per well diluted in 500 μL of serum-free DMEM was added to the top of the chambers, with 500 μL of media containing 10% FBS in the bottom chambers. Cells were incubated for 24 hours, after which the tops of the chambers were scrubbed with sterile cotton swabs, and the bottoms were fixed in 4% paraformaldehyde (PFA) for 15 minutes, permeabilized in 0.2% Triton X-100 in PBS for 30 minutes, blocked in 5% BSA in PBS for 30 minutes, and stained with Hoechst dye (1:2,000 dilution) for one hour. Images were captured at 40X total magnification via epifluorescence microscopy, and cells were counted using ImageJ. Invasion was calculated as the proportion of cells migrating through Matrigel-coated chambers divided by the number of cells migrating through uncoated chambers.

Scratch wound migration assays were performed in 96-well plates for 143B cells and in 24-well plates for RD cells. 143B cells were seeded at 20,000 cells per well and incubated overnight, after which scratches were made in the wells using a WoundMaker (Essen Biosciences). For RD cells, 200,000 cells per well were seeded and incubated overnight. Scratches were made using a 20-200 μl pipette tip. For both cell types, wells were washed with PBS, and replaced with fully supplemented DMEM. Images were captured every two hours using the IncuCyte Zoom Live Cell Imaging System. Scratch wound migration of 143B cells was analyzed using the migration analysis module in the IncuCyte Zoom software, and migration of RD cells was analyzed using ImageJ.

For monolayer proliferation assays, 25,000 cells per well were plated in 500 μL of DMEM in 24-well plates, and percent confluence was analyzed using the IncuCyte Zoom software, with imaging at two hour intervals at 100X total magnification.

Soft agar anchorage-independent growth assays were performed as previously described (30). Briefly, 25,000 cells were embedded in a 0.3% agarose layer atop a bottom layer of 0.5% agarose and cultured for approximately three weeks in fully supplemented DMEM in 6-well plates. Media was replenished weekly, after which colonies were stained with Nitrotetrazoleum Blue overnight, plates were imaged, and colonies counted using ImageJ.

For spheroid growth assays, cells were scraped in fully supplemented DMEM, and 1,000 cells/well were seeded in ultra-low attachment plates (Costar, cat. #7007). Spheroids were quantified by manual counting after approximately 2 weeks in culture.

For cell-cell adhesion assays, 12-well plates were incubated overnight with 0.5%BSA/Fraction V in CMFS buffer at 4°C. CMFS buffer contains 4 g NaCl, 0.2g KCl, 0.03g Na_2_HPO_4_, and 0.5 g glucose diluted in 500 ml PBS (-/-). Plates were washed with CMFS buffer and 200,000 cells were immediately seeded in 1mL of media. Plates were incubated on an orbital shaker at 100 rpm at 37°C for 3 hours, followed by the addition of 1.5 mL of 4% PFA and incubated at room temperature for 30 minutes. Cells aggregates were immediately quantified in size categories using ImageJ.

### Statistical analyses

All data were analyzed for statistically significant differences using the Student’s t-test (for two comparisons) or analysis of variance with Tukey’s post-hoc correction (for multiple comparisons) in JMP Pro 13. Any p-value < 0.05 was considered statistically significant.

### Quantitative modeling for simulating the mechanical effects of E-cadherin

We use a two-dimensional subcellular elements model (31–33) to simulate the effects of different levels of E-cadherin, where each cell is represented by two subcellular elements, the front and rear element (inserted cartoon in Fig. 7B (32)). Each subcellular element is self-propelled with a self-propulsion force m, which balances the intracellular contraction f_contr_ between the front and rear element. We use a long range attractive and short range repulsive intercellular force f_rep/adh_ to model cell-cell adhesion (E-cadherin level) and volume exclusion, respectively. There is a friction between each subcellular element and the substrate with a constant coefficient ξ. The velocity and position for each subcellular element is updated by: v × 1/ξ (m + f_contr_ + f_rep/adh_), and x × vdt. In our simulation, cells can divide with a probability based on its size (32). Once the cell length exceeds a certain threshold, it divides at a given probability. Upon division, two new subcellular elements are inserted, forming two new cells. This process simulates the cell cycle: the expansion of cell culture is driven by contact inhibition of locomotion, and cells are elongated during the expansion.

### Quantitative mathematical modeling of E-cadherin regulatory network in mediating anchorage independence

A mathematical model representing the interactions among E-cadherin, TBX2, NRAGE, and their effects on anchorage independence was constructed. The set of coupled ordinary differential equations (ODEs) representing the dynamics of these molecular species was simulated in MATLAB (Mathworks Inc.). The details of model construction and parameters used is given in supplemental methods.

## Results

### Aberrant E-cadherin expression in sarcomas is prognostic for improved clinical outcomes

Our previous work found that phenotypic plasticity in sarcomas leads to a more epithelial-like phenotype (2). This transition to a more epithelial gene expression program has prognostic relevance: Compared to patients whose sarcomas are more strongly mesenchymal-like, patients with epithelial-like sarcomas have improved overall survival (2). One of the hallmarks of this mesenchymal-epithelial transition (MET) is upregulation of the epithelial cell-cell adhesion molecule, E-cadherin. To further understand the prognostic relevance of E-cadherin in sarcomas, we stratified osteosarcoma patients into E-cadherin^high^ and E-cadherin^low^ groups using publicly-available RNA-Seq data (27). E-cadherin^high^ patients had improved metastasis-free survival (Figure 1A) and overall survival compared to E-cadherin^low^ patients (Figure 1B). Likewise, soft tissue sarcoma (STS) patients from The Cancer Genome Atlas with higher E-cadherin mRNA trended toward better overall survival than patients with lower E-cadherin (Figure 1C) while patients with higher E-cadherin protein had significantly improved overall survival compared to patients with lower E-cadherin protein levels (Figure 1D). These analyses suggest that E-cadherin has prognostic utility in sarcomas.

### Ectopic E-cadherin expression in sarcomas does not promote MET

While it is clear that E-cadherin has prognostic relevance in sarcomas, the underlying molecular mechanism(s) and cellular phenotypes by which E-cadherin contributes to a less aggressive tumor remain unknown. Given the important role of E-cadherin as a mediator of the epithelial phenotype, we hypothesized that E-cadherin may promote an MET-like phenotype in sarcomas, including suppression of migration and invasion pathways. Consistent with this hypothesis, data from The Cancer Genome Atlas indicate E-cadherin expression in sarcomas is inversely correlated with an EMT signature; sarcomas with a more mesenchymal-like gene expression program have lower E-cadherin while those with a more epithelial-like gene expression program have higher E-cadherin (Figure 2A). To test the hypothesis that E-cadherin inhibits sarcoma aggression by promoting an MET-like phenotypic switch, we ectopically expressed E-cadherin in RD human rhabdomyosarcoma and 143B human osteosarcoma cells and assayed for changes in cell growth. We found E-cadherin expression had no effect on the growth of cells in monolayer (**Supplementary Figure 1B, 1D**). We next assayed for changes in EMT biomarkers (Snail, Slug, Twist, Zeb1, Vimentin) by qRT-PCR. With the exception of a statistically significant two-fold reduction in Slug in 143B cells, E-cadherin had no significant effect on EMT markers (Figure 2B, **Supplementary Figure 2A**). Consistent with these results, E-cadherin expression did not alter scratch wound migration (Figure 2C) or invasion in 143B cells (Figure 2D) or in RD cells (**Supplementary Figure 2B-C**). We next asked if E-cadherin status changed the overall EMT score using our previously-derived scoring metric for epithelial, mesenchymal and hybrid epithelial/mesenchymal states (George, Jolly et al. 2017). Using empirical probably density functions, we calculated EMT scores for E-cadherin^high^ vs. E-cadherin^low^ expressing osteosarcomas. The scoring metric was strongly mesenchymal for all sarcomas analyzed, regardless of E-cadherin status, suggesting that E-cadherin was not capable of lineage reprogramming toward an epithelial-like state in sarcomas (Figure 2E). Visualization of the EMT score distribution shows only a slight increase in the number of samples predicted as epithelial or hybrid E/M in cases of E-cadherin^high^ samples relative to the E-cadherin^low^ samples (Figure 2E). Together, using both computational and experimental approaches, our results suggest that E-cadherin overexpression need not be sufficient to induce MET in sarcomas, and that E-cadherin acts independently of migration/invasion programs to inhibit sarcoma aggressiveness.

### E-cadherin suppresses anchorage-independent growth in sarcomas

To explore alternative mechanisms by which E-cadherin may impact the aggressiveness of sarcoma tumors, we analyzed the effect of E-cadherin on anchorage-independent growth. Given the role of E-cadherin as a cell adhesion molecule, we posited that E-cadherin may alter the anchorage-independent growth phenotype of sarcoma cells. To test this hypothesis, cells expressing an empty vector (EV) or E-cadherin were cultured in soft agar to simulate anchorage-independent growth. Interestingly, E-cadherin expression significantly inhibited the number of colonies formed in soft agar growth assays in 143B cells (Figure 3A). This effect was independent of differences in monolayer growth, as E-cadherin expressing cells showed no difference in 2D growth (**Supplementary Figure 1B, D**). We observed similar results in a second sarcoma cell line, RD cells. Ectopic expression of E-cadherin did not induce MET by gene expression, migration, or invasion (**Supplementary Figure 2A-C**). However, consistent with 143B cells, E-cadherin overexpression inhibited anchorage independent growth (**Supplementary Figure 2D, E**). We next measured spheroid formation to assess the impact of E-cadherin on anchorage-independent growth and morphology. For these experiments, we noted that, despite consistent downregulation of soft agar growth in both RD and 143B cells, E-cadherin was not expressed at the cell membrane of RD cells (**Supplementary Figure 1A**), while E-cadherin was localized at the membrane of 143B cells (**Supplementary Figure 1C**). Because membrane E-cadherin is the most typically-characterized aspect of E-cadherin function, we focused on 143B cells for which E-cadherin was correctly localized to the cell membrane. We also included U2OS cells as a second model for which E-cadherin was localized to the cell membrane (**Supplementary Figure 1E**). Consistent with our result in soft agar, E-cadherin expressing spheroids formed as tight aggregates and significantly reduced the size of spheroid formation in both 143B and U2OS cells (Figure 3B, C). These results indicate that sarcomas with E-cadherin expression have suppressed anchorage-independent growth as indicated by their deficient ability to grow suspended in soft agar.

### E-cadherin-mediated sensitivity to anchorage-independent growth acts, in part, through CREB

To determine the signaling pathway(s) through which E-cadherin inhibits anchorage-independent growth in sarcomas, we performed phospho-kinase arrays on 143B osteosarcoma cells with and without E-cadherin expression. The phospho-kinase arrays contain antibodies against 43 different kinases (R&D Systems). From these arrays, we identified phospho (p)-CREB as consistently downregulated in E-cadherin expressing 143B cells as compared to 143B cells expressing an empty vector lacking E-cadherin (Figure 4A). The inhibition of phospho- and total CREB in E-cadherin expressing 143B cells was validated by enzyme linked immunosorbent assays (ELISAs) and western blotting (Figure 4B, C). Consistent with this, we also observed lower levels of phospho- and total CREB in Abrams canine osteosarcoma cells expressing E-cadherin as compared to cells expressing an empty vector lacking E-cadherin (Figure 4D).

We next asked if E-cadherin-mediated CREB inhibition was responsible for the reductions in anchorage-independent growth capacity of E-cadherin expressing cells. To do this, we knocked down CREB using two independent siRNAs and verified knockdown at the mRNA and protein levels in 143B cells, with siRNA_5 providing a stronger knockdown (Figure 4E, F). We noted a consistent reduction in soft agar growth with both CREB siRNAs compared to a nonsilencing siRNA (Figure 4G); however, there was only a significant inhibition of soft agar growth in cells treated with the most effective siRNA_5 (Figure 4G). Importantly, CREB knockdown had no effect on E-cadherin expression (**Supplementary Figure 3A**). We conclude from these data that E-cadherin inhibition of anchorage-independent growth is mediated through CREB, and downregulation of CREB partially phenocopies E-cadherin expression, although other factors are likely at play in driving anchorage-independent growth in sarcomas.

During our analysis of MET biomarkers and phenotypes in E-cadherin expressing cells, we noted that groups of E-cadherin+ 143B cells lost cell surface N-cadherin expression (**Supplementary Figure 4A**). Interestingly, the reduction in cell surface N-cadherin was not due to changes in N-cadherin expression at either the mRNA or protein levels (**Supplementary Figure 4B, C**). Cadherin switching has been observed in carcinomas in which an E-cadherin to N-cadherin switch is prognostic for poorer clinical outcomes (34–36). In addition, N-cadherin has been shown to promote anchorage-independent growth in carcinomas (37). Thus, we tested if a reduction of cell surface N-cadherin was responsible for the downregulation of anchorage-independent growth in E-cadherin+ cells. Knockdown of N-cadherin with two independent siRNAs (**Supplementary Figure 4D**) had no effect on 143B colony formation in soft agar (**Supplementary Figure 4E**). Likewise, N-cadherin knockdown did not alter levels of p-CREB protein, as determined by p-CREB ELISAs and western blotting (**Supplementary Figure 4F, G**). These results suggest that E-cadherin inhibits anchorage-independent growth through a pathway that is independent of N-cadherin.

### An E-cadherin/TBX2/CREB axis modulates anchorage-independent growth of sarcomas

To elucidate further the signaling axis that modulates E-cadherin/CREB-mediated suppression of anchorage-independent growth, we investigated additional signaling axes that may be involved with E-cadherin. One of these axes, TBX2, is a transcription factor that can directly repress E-cadherin (38), can be modulated by CREB (39), and is overexpressed in rhabdomyosarcoma (40,41). To determine if E-cadherin regulates TBX2 in sarcomas, we measured levels of TBX2 in E-cadherin expressing 143B cells. TBX2 mRNA was significantly downregulated in E-cadherin+ 143B cells (Figure 5A). Similarly, U2OS cells ectopically expressing E-cadherin (**Supplementary Figure 1E**) had significantly lower levels of TBX2 mRNA (Figure 5B). To further delineate the directionality of the E-cadherin/CREB/TBX2 signaling network, we performed siRNA-mediated CREB knockdown and measured TBX2 mRNA levels. Interestingly, CREB knockdown had no effect on TBX2 expression (Figure 5C, **Supplementary Figure 3B**), suggesting that CREB is downstream or independent of TBX2 in this gene regulatory network. Consistent with these data, TBX2 knockdown (Figure 5D) led to significant reductions in CREB mRNA (Figure 5E) and protein (Figure 5F). Taken together, these results suggest that E-cadherin inhibits TBX2, which leads to CREB downregulation and suppression of anchorage-independent growth.

### Mathematical modeling of E-cadherin/TBX2 axis regulating anchorage-independent growth

By integrating our quantitative RT-PCR result showing E-cadherin-mediated suppression of TBX2 mRNA with previous reports suggesting E-cadherin to be a transcriptional target of TBX2 (38), and regulation of anchorage-independent growth by E-cadherin and/or TBX2 (40,42,43), we constructed a mechanism-based mathematical model to decode the dynamics of the E-cadherin/TBX2 axis in mediating anchorage-independent growth (Figure 6A). This model includes the following interactions: (i) mutual repression between E-cadherin and TBX2, (ii) inhibition of NRAGE (neurotrophin receptor-interacting melanoma antigen) by E-cadherin (Kumar et al., 2011), and (iii) inhibition of anchorage-independent growth by the TBX2 complex, which is promoted in the presence of NRAGE (42). Such interconnected regulatory loops can often obviate an intuitive understanding of the emergent outcomes of these interactions (44). Thus, mechanism-based mathematical models can be a powerful tool for both helping explain previous experimental observations, and for making novel predictions to guide future experimental design (45).

Our model predicted that TBX2 knockdown would restrict anchorage-independent growth in a similar way as that by E-cadherin overexpression (Figure 6B). To test this prediction, we used two independent siRNA to TBX2 to knockdown expression and measured anchorage independent growth. Consistent with our prediction, TBX2 knockdown using two independent siRNAs significantly abrogated colony growth in 143B cells (Figure 6C). Further, the model predicted that knockdown of TBX2 under ectopic expression of E-cadherin accentuates sensitivity to anchorage independence (Figure 6B). However, TBX2 knockdown had no effect on spheroid formation, when E-cadherin was overexpressed in U2OS cells (Figure 6D).

The inconsistency in model predictions vs. experimental observations drew our attention to multiple underlying assumptions of the mathematical model – (a) TBX2-mediated inhibition of E-cadherin affects both mRNA and protein levels of E-cadherin, (b) endogenous production of E-cadherin is enough to inhibit TBX2. The first assumption is based on observations in breast epithelial cells MCF10A (38); similarly, the second assumption also is likely to be more typical of carcinomas than of sarcomas. Given that our model was constructed using data exclusively taken from carcinomas, we reasoned that the discrepancy between the model and the experimental validation may be due to differences in regulatory networks between carcinomas and sarcomas. Therefore, to determine if similar regulatory networks between TBX2 and E-cadherin exist in sarcomas, we knocked down TBX2 with two independent siRNAs in U2OS cells and measured E-cadherin mRNA and protein levels. TBX2 knockdown led to a modest, albeit significant, two-to four-fold upregulation of E-cadherin mRNA levels (Figure 6E). More importantly, this modest change in expression in a cell line with relatively low levels of E-cadherin mRNA expression at baseline was not enough of an increase in mRNA to show a change at the protein level (Figure 6F). The relative lack of E-cadherin re-activation upon TBX2 knockdown led us to hypothesize that sarcoma cells had alterations in the promoter of E-cadherin that prevented robust increases in expression upon TBX2 downregulation. Consistent with this hypothesis, analysis of sarcoma tumors from The Cancer Genome Atlas (TCGA) revealed that the E-cadherin promoter was significantly more methylated in sarcomas as compared to that in carcinomas (Figure 6G). Moreover, the degree of E-cadherin promoter methylation was lowest in synovial sarcoma (Figure 6H), which corresponded with a relative increase in E-cadherin expression (Figure 6I). Interestingly, synovial sarcoma often displays epithelioid histological features, including E-cadherin upregulation. This suggests that one potential difference between the E-cadherin/TBX2 axis in sarcomas vs. carcinomas is the ability of TBX2 to access the E-cadherin promoter, which may impact the mathematical models of networks based from largely studies in carcinoma models. Incorporating this new experimental data, we revised our mathematical model to include a weaker inhibition of E-cadherin by TBX2. The revised model predicted that, consistent with our experimental results, the effect of TBX2 knockdown on anchorage-independent growth in the context of E-cadherin overexpression was reduced (**Supplementary Figure 5**).

### A mechanical model relates E-cadherin mediated cell-cell adhesion to spheroid formation

Our experimental results suggest E-cadherin inhibits soft agar colony formation mediated by suppression of CREB and TBX2 in sarcoma cells. Interestingly, E-cadherin expression alone is sufficient to suppress the size of spheroid formation while TBX2 has no effect on spheroid growth (Figure 6D). The differences in response between these phenotypic assays suggest that E-cadherin may act through different molecular mechanisms to suppress anchorage-independent growth and spheroid growth. Given the well-described role for E-cadherin as an adhesion molecule, we hypothesized that E-cadherin can regulate spheroid formation through its mechanical role as a regulator of cell-cell adhesion. Indeed, E-cadherin mediated cell-cell adhesion has been proposed to be crucial in mediating formation of cohesive multicellular units such as tumor emboli and invasive collective migration in breast cancer (19,46–48). Consistent with this, U2OS cells overexpressing E-cadherin formed tight multicellular clusters at 24 hours of spheroid formation as compared to control cells (Figure 7A).

To quantitatively capture the relationship between cell-cell adhesion and spheroid size, we developed a two-dimensional mechanical model of sphere formation. This model represents each cell by two subcellular elements connected to each other with a contractile force, and incorporates various interconnected biomechanical aspects of cellular aggregation, such as intracellular contraction, intercellular adhesion, cell proliferation, self-propulsion forces of the cells, and friction between the cells and the substrate (31–33). We initialized our simulations with a seed colony of 233 cells arranged as a two-dimensional circular sheet, and simulated the colony growth for varying levels of intercellular adhesion, while keeping all other parameters the same. The intercellular interaction is repulsive at short distances, attractive at longer distances and becomes zero further away, which models the volume exclusion and cell-cell adhesion. For different levels of adhesion, the attractive part of the intercellular force follows different curves (Figure 7B). This mechanical model predicts a decrease in final spheroid size and the total number of cells with an increase in E-cadherin levels (Figure 7C), and a concomitant increase in cell density (**Supplementary Figure 6**). Thus, the model predicts that cell-cell adhesion can regulate colony size through mechanical factors.

To test if E-cadherin modulates cell-cell adhesion in sarcomas, we quantified the numbers of single cells and cellular aggregates (two or more cells clumped together) using cellcell adhesion assays for both 143B and U2OS cells upon ectopic expression of an empty vector or E-cadherin. Consistent with our model, we observed that E-cadherin overexpression drives a significant enrichment of multicellular aggregates in both 143B and U2OS cells (Figure 7D-E), suggesting that E-cadherin levels can play a fundamental role in enabling cell-cell adhesion in sarcomas. Together, our results integrating gene regulatory network models, mechanical models, and experimental validations indicate that E-cadherin plays a dual role in suppressing sarcoma aggression that is independent of EMT through both signaling-related inhibition of anchorage-independent growth and mechanics-based inhibition of spheroid formation.

## Discussion

Our integrated computational-experimental study illustrates that E-cadherin inhibits sarcoma aggressiveness in at least the following two ways: first, by suppressing anchorage-independent growth through a CREB/TBX2 signaling axis; and second, by increasing cell-cell adhesion and restricting spheroid growth. Anchorage independent growth is a hallmark of cancer and a contributor to metastatic growth (50,51). Although normal cells require attachment to the ECM to receive cell survival signals, cancer cells can overcome the death signals that are driven by loss of cell-ECM attachment (49). This can lead to anchorage-independent survival of otherwise adherent cells, including at sites where a unique ECM composition prevents cellular attachment (49). Interestingly, anchorage-independent growth has often been studied in the context of EMT in multiple epithelial cancers (52–54); however, ours is among the minority of studies connecting E-cadherin with anchorage independence in sarcomas. Indeed, our results pinpoint an E-cadherin/p-CREB/TBX2 axis that mediates anchorage-independent growth in sarcoma cells. This signaling axis is independent of MET, as neither ectopic E-cadherin expression nor TBX2 knockdown had any effect on MET markers in 143B or RD cells. This observation reinforces previous results that overexpression of transcription factor GRHL2 – an activator of E-cadherin – was sufficient to induce MET in breast cancer cells having undergone EMT, but not in sarcoma (2,55). Highly methylated promoters of both E-cadherin (Figure 6G) and GRHL2 (2) in sarcomas relative to carcinomas, as observed from analysis of TCGA data, can potentially explain this difference. In this scenario, epigenetic silencing of epithelial promoters would prevent lineage reprogramming and MET. However, it is possible that some sarcoma subtypes are more likely to undergo MET; for instance, E-cadherin expression (**Supplementary Figure 1A**), and an epithelial-like phenotype in general (56), appears to be more frequently upregulated in certain soft tissue sarcoma subtypes, such as synovial sarcomas, malignant peripheral nerve sheath tumors, and leiomyosarcomas (Figure 6I).

While E-cadherin inhibits anchorage-independent growth through a signaling axis, the role of E-cadherin in suppressing spheroid growth is mediated, at least in part, through its mechanical activity of increasing cell-cell adhesion and cellular aggregation (Figure 7). This observation is consistent with previous reports in which knock down of E-cadherin led to less cell-cell adhesion and lower multicellular aggregation in colorectal cancer cells (57). However, unlike our study, this study in colorectal cancer also found that E-cadherin loss led to increased migration and invasion. Yet, despite the fact that E-cadherin did not change migration or invasion in sarcomas, E-cadherin may still inhibit invasiveness of some tumors not by maintaining an epithelial state, but by increasing cell-cell adhesion. In this way, E-cadherin would indirectly reduce invasion by preventing cells from detaching from the bulk tumor without changing the EMT factors involved in migration and/or invasion.

The signaling and mechanical modes of action we observed in suppressing anchorage-independent growth and spheroid formation, respectively, need not be independent of each other, especially given the interconnections among mechanosensitive and biochemical signaling pathways. For instance, Notch signaling can be mechanosensitive (58), give rise to multiple multicellular patterns of cells in varying phenotypic states (59,60), and be involved in anoikis (61). Similarly, E-cadherin can sense and provide mechanical cues for epithelial homeostasis and collective cell migration (62–64). Future studies will be aimed at elucidating the crosstalk between inter-cellular mechano-sensitive and intra-cellular p-CREB/TBX2 modules to yield new insights into the mechanisms underlying non-cell autonomous control of anoikis, as recently observed (65). Moreover, it would be intriguing to compare the results for soft agar colony growth and spheroid formation assays for RD cells where E-cadherin exerts an effect on anchorage-independent growth in the absence of membrane localization.

Overall, our results provide a mechanistic basis for the observation that E-cadherin is a prognostic biomarker for improved clinical outcomes in sarcomas. Although E-cadherin upregulation in the clinical setting is likely to be a harbinger of a broader change in EMT/MET programs, our data argue that E-cadherin upregulation also has important effects on sarcoma aggressiveness that are independent of MET induction. Together, these results indicate that targeting the EMT/MET axis in sarcomas may have multiple benefits, including downregulation of stemness, migration, invasion, and anchorage-independent growth pathways. One way this might be achieved is through epigenetic reprogramming. Indeed, multiple reports in carcinomas suggest that treatment with epigenetic modifiers, including histone deacetylases inhibitors, can upregulate E-cadherin (66–69). Consistent with this hypothesis, when anchorage-dependent and anchorage-independent subpopulations of osteosarcoma cells are compared, the anchorage-independent subpopulations are significantly more susceptible to epigenetic modifying therapies (70). A similar strategy could be used to drive sarcomas toward a more epithelial-like phenotype and lower their aggressiveness by targeting many aggressive phenotypes simultaneously. Such combinatorial therapies may limit the adaptive phenotypic plasticity of cancer cells – a crucial determinant of therapeutic resistance (71).

## Supporting information

Supplementary Infomation

## Acknowledgments

The results published here are in part based upon data generated by the TCGA Research Network: http://cancergenome.nih.gov. The authors acknowledge the Duke University Cell Culture Facility and the Duke Cancer Institute Flow Cytometry Shared Resource. HL was supported by NSF Center for Theoretical Biological Physics (NSF PHY-1427654) and NSF DMS-1361411, and as a CPRIT (Cancer Prevention and Research Institute of Texas) Scholar in Cancer Research of the State of Texas at Rice University. MKJ received a training fellowship from the Gulf Coast Consortia on the Computational Cancer Biology Training Program (CPRIT grant no. RP170593). KEW was supported by the NIH F32 CA192630. JAS wishes to acknowledge support from Meg and Bill Lindenberger, the Paul and Shirley Friedland Fund, the Triangle Center for Evolutionary Medicine, and funds raised in memory of Muriel E. Rudershausen (riding4research.org).

---

**Figure 1.**
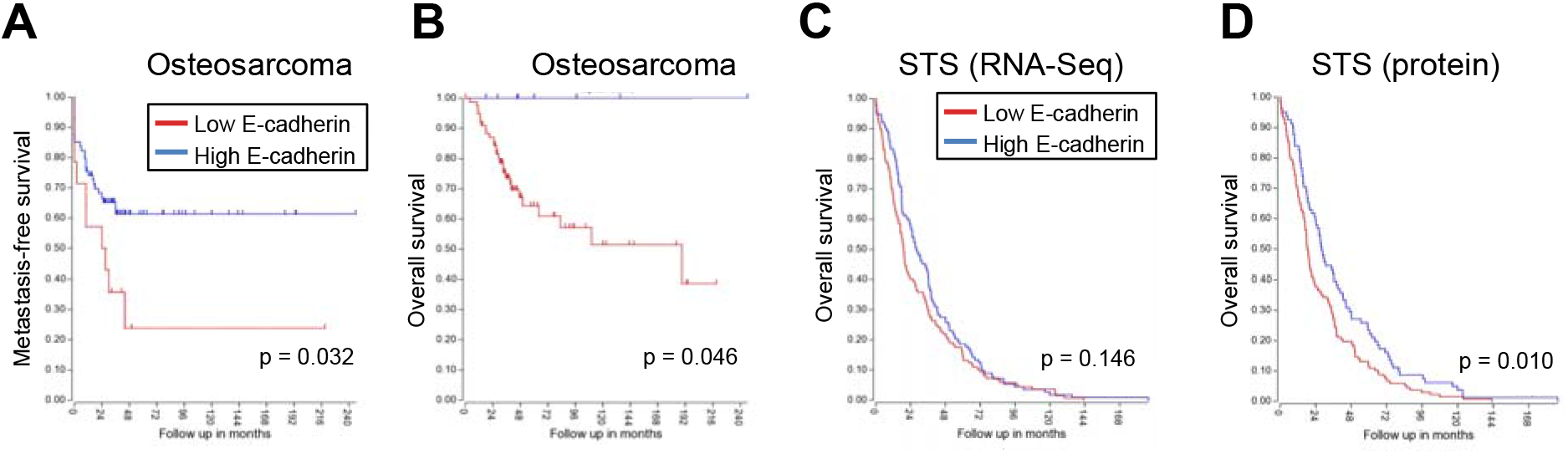
E-cadherin upregulation is prognostic for improved outcomes in sarcoma. **A-B**. Osteosarcomas with elevated E-cadherin have better metastasis-free survival **(A)** and overall survival **(B)** as compared to tumors with low/no E-cadherin expression. **C-D**. Soft tissue sarcomas (STS) from The Cancer Genome Atlas with higher E-cadherin mRNA **(C)** and protein expression **(D)** have improved overall survival as compared to tumors with low or no E-cadherin.

**Figure 2.**
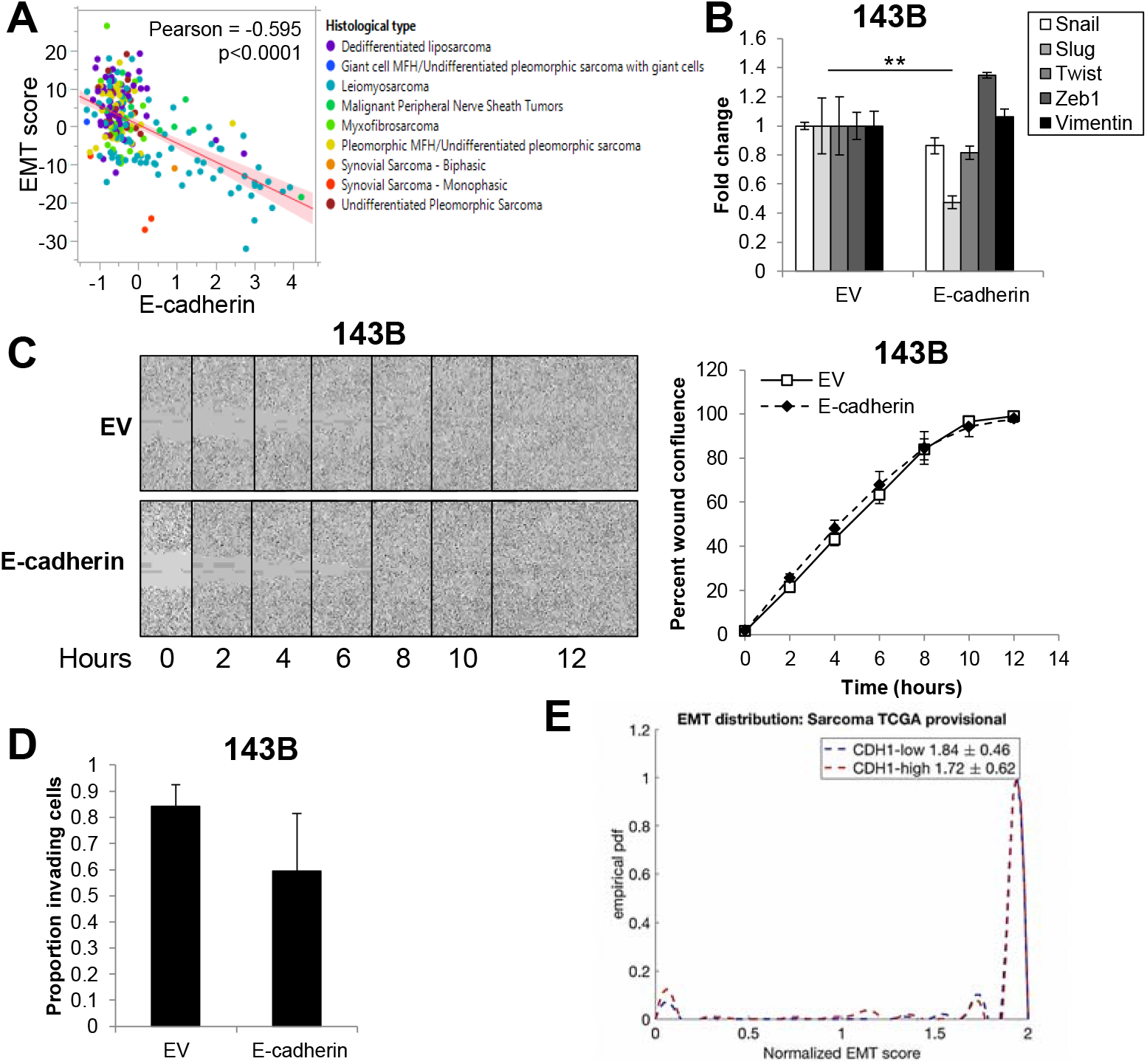
Ectopic E-cadherin expression in sarcoma cells does not alter EMT. **A.** E-cadherin expression significantly correlates with the EMT score of sarcoma tumors in The Cancer Genome Atlas. **B.** Ectopic expression of E-cadherin in 143B human osteosarcoma cells has no influence on mesenchymal markers (Snail, Slug, Twist, Zeb1, Vimentin). **C.** E-cadherin expression has no effect on migration of 143B cells. Images (left) were collected every two hours and quantified (right) using the IncuCyte Zoom system. **D.** E-cadherin expression does not change invasion in 143B cells. **E.** Using mRNA expression of CDH1 (E-cadherin), empirical probably density functions of EMT scores for CDH1-high (red) and CDH1-low (blue) TCGA sarcoma sample were constructed by interpolation of the EMT score histogram (E<0.5, 0.5≤E/M≤1.5, M>1.5). EMT score distribution showed no significant difference when separated for E-cadherin^high^ vs. E-cadherin^low^ expression from analysis of publicly-available sarcoma data sets.

**Figure 3.**
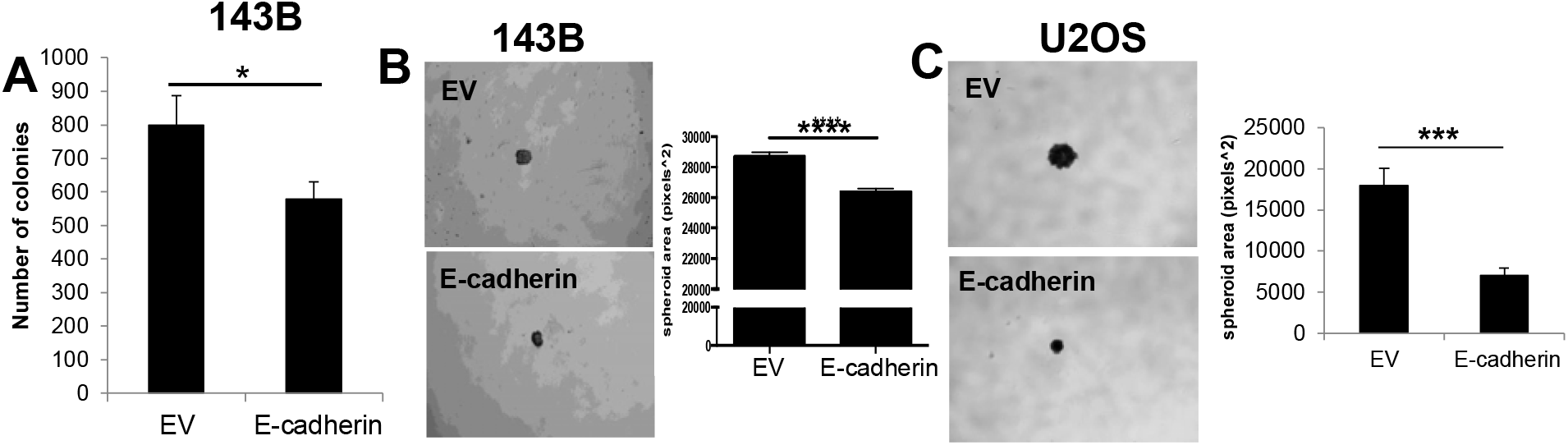
E-cadherin inhibits anchorage-independent growth of sarcomas. **A.** Anchorage-independent growth of 143B cells expressing E-cadherin was significantly inhibited. **B-C.** E-cadherin expression leads to reduces spheroid size in **B.** 143B and **C.** U2OS cells.

**Figure 4.**
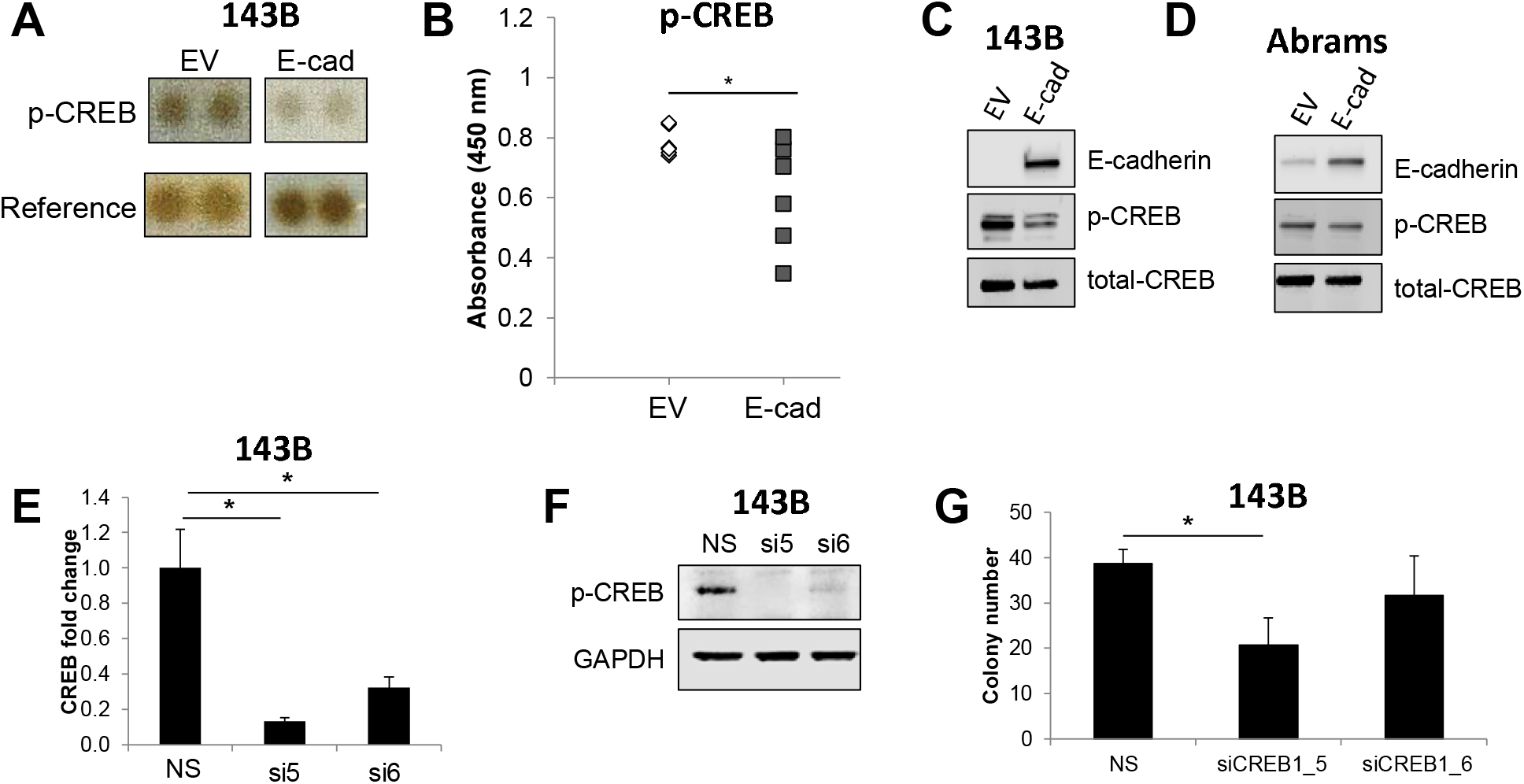
Ectopic expression of E-cadherin in sarcoma cells inhibits phospho-CREB levels. **A.** A phospho-kinase array revealed that E-cadherin led to downregulation of phospho-CREB. **B-C**. E-cadherin-mediated phospho-CREB inhibition was verified by B. ELISAs and **C.** western blotting. **D.** Abrams canine osteosarcoma cells exhibited reduced phospho-CREB in E-cadherin over-expressing cells. **E.** QRT-PCR confirmed knockdown of CREB with two independent siRNAs. **F.** Western blotting to confirm knockdown of CREB in 143B cells. **G.** CREB knockdown led to a modest downregulation of 143B colony growth in soft agar.

**Figure 5.**
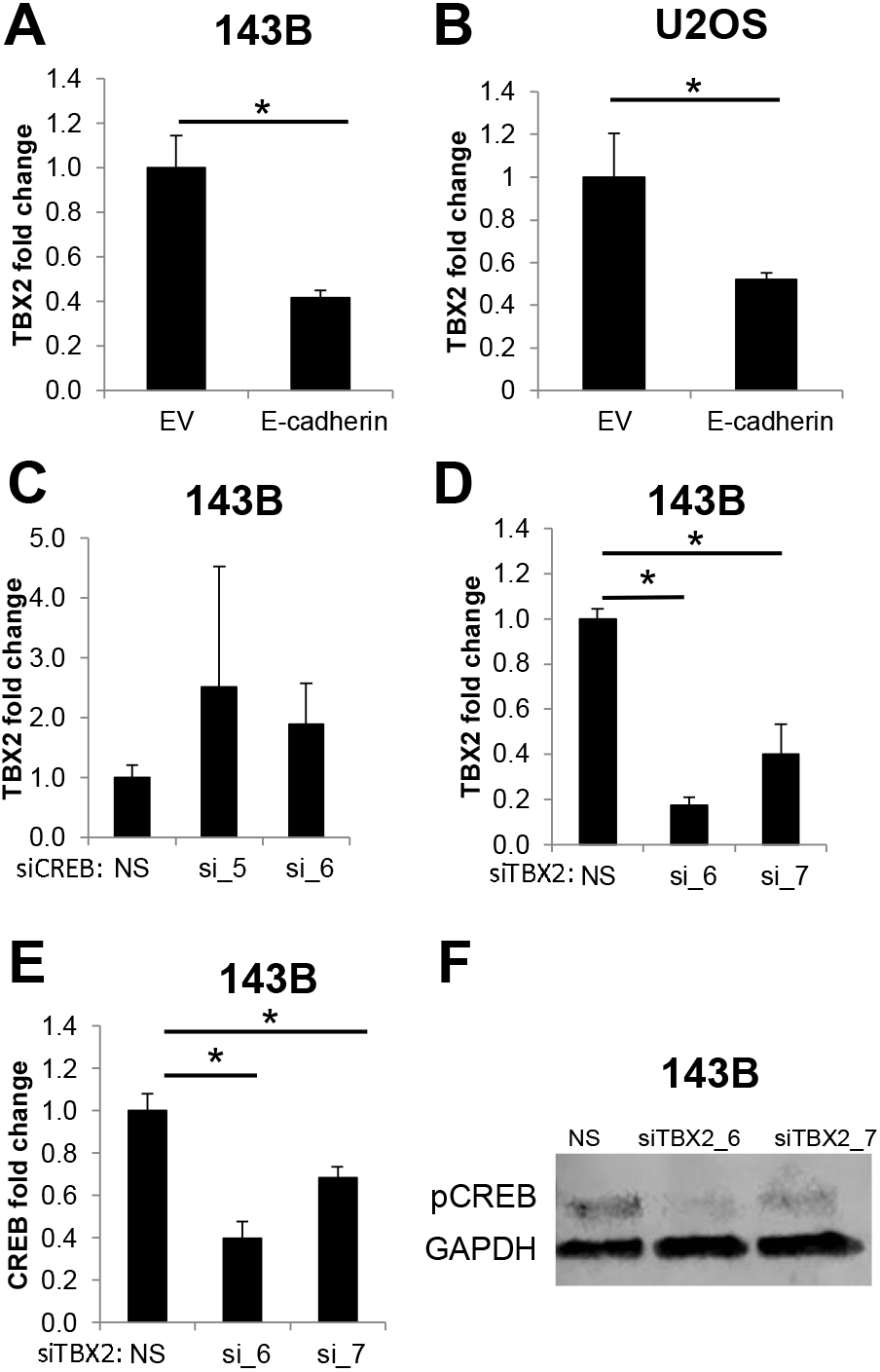
TBX2 knockdown phenocopies E-cadherin-mediated CREB inhibition. **A-B.** TBX2 mRNA is downregulated in E-cadherin-expressing **A.** 143B cells and B. U2OS cells. **C.** CREB knockdown with two independent siRNAs had no effect on TBX2 mRNA. **D-F.** Conversely, TBX2 knockdown, verified in **D.** led to a significant reduction in CREB mRNA **(E)** and protein level **(F)**.

**Figure 6.**
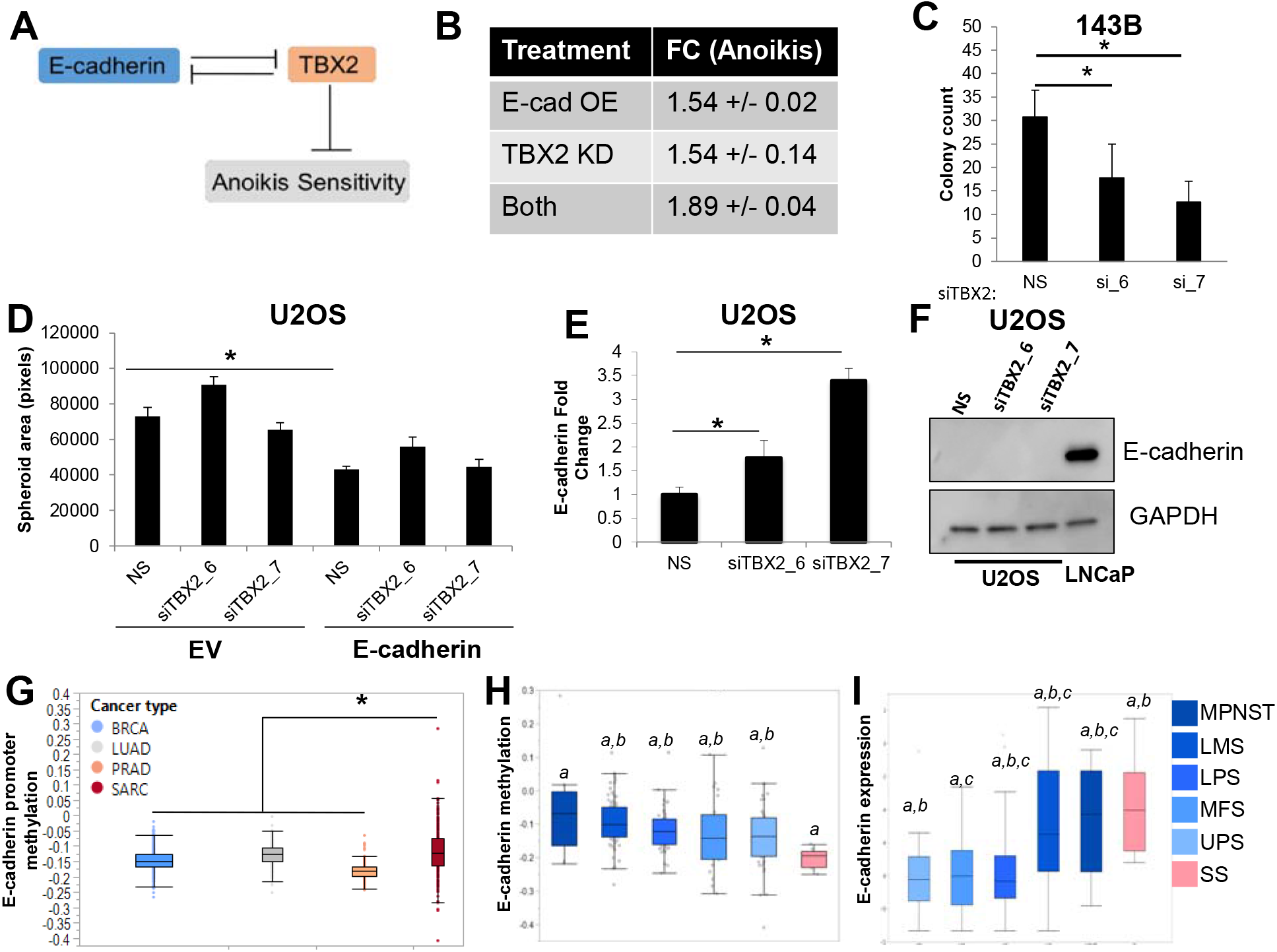
E-cad/TBX2 interplay mediates anchorage-independent growth. **A.** Intracellular regulatory circuit modulating anchorage-independent growth. **B.** Mathematical model predicts similar effects of Ecad-OE and TBX2-KD on anchorage-independent growth. **C.** TBX2 knockdown using two independent siRNAs inhibited anchorage-independent growth of 143B cells. **D.** TBX2 downregulation did not affect sphere formation in the presence of ectopic E-cadherin expression. **E.** TBX2 knockdown upregulates E-cadherin mRNA. **F.** Western blot analysis of U2OS cells upon TBX2 knockdown indicates E-cadherin protein is not upregulated by loss of TBX2. Prostate cancer (PC) cell line, LNCaP, was used as a positive control for E-cadherin expression. **G.** The E-cadherin promoter is differentially methylated in sarcomas as compared to carcinomas. **H.** Synovial sarcomas, which often display epithelioid histopathological features, have the lowest levels of E-cadherin promoter methylation of all sarcoma subtypes. **I.** Among soft tissue sarcoma histological subtypes E-cadherin mRNA expression is highest in synovial sarcomas. Letters indicate statistically significant associations between groups. Undifferentiated pleomorphic sarcoma (UPS), liposarcoma (LPS), leiomyosarcoma (LMS), synovial sarcoma (SS), malignant peripheral nerve sheath tumors (MPNSTs), and Myxofibrosarcoma (MFS).

**Figure 7.**
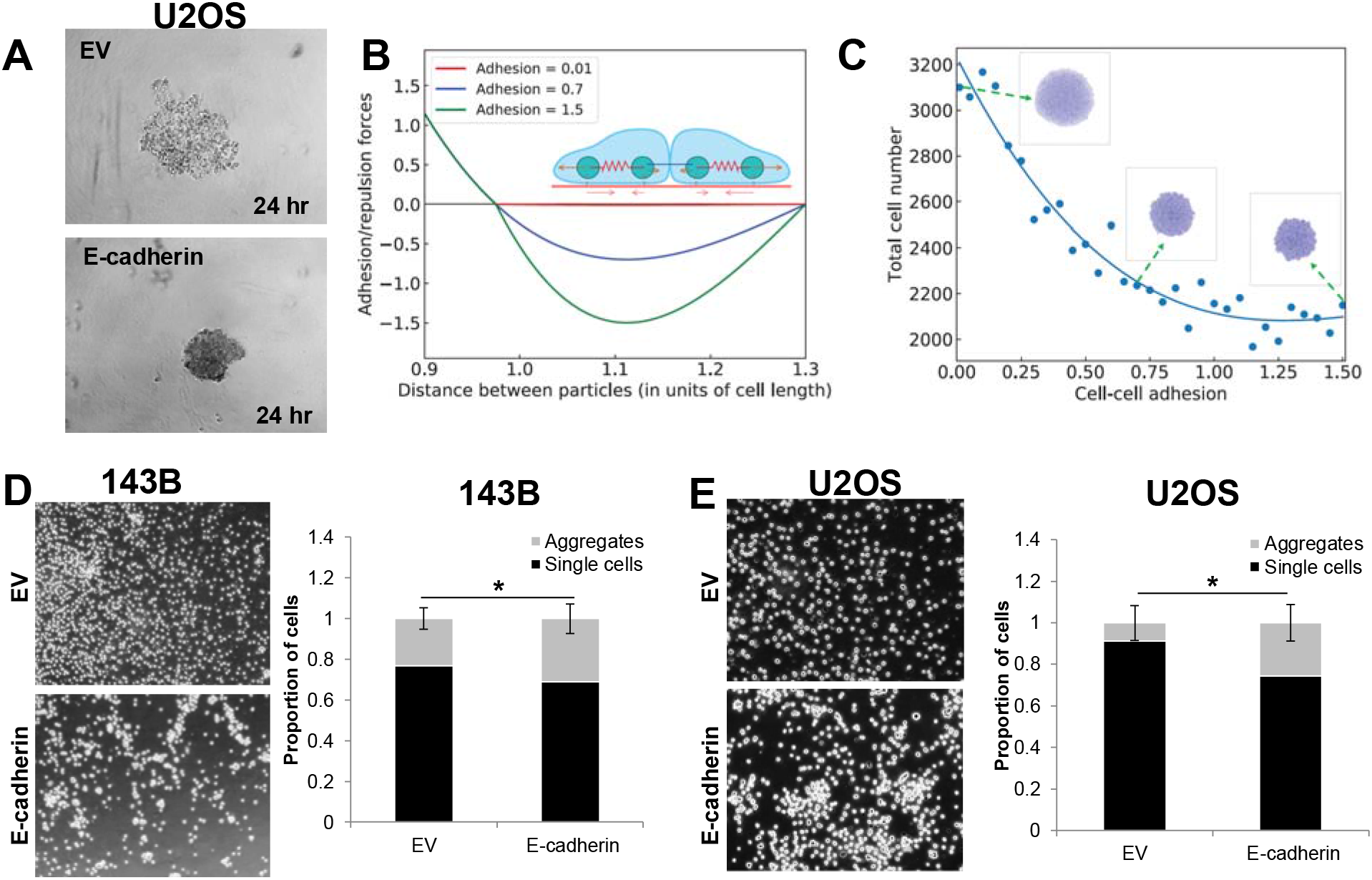
E-cadherin inhibits spheroid formation by increased cell-cell adhesion. **A.** E-cadherin causes tighter clustering of cells during spheroid formation. **B.** A mechanical model illustrates the relationship between particle distance to adhesion. **C.** A mechanical model predicts E-cadherin drives down spheroid size through an increase in cell-cell adhesion. **D-E.** As predicted in the model, ectopic E-cadherin expression increases cell-cell adhesion in 143B **(D)** and U2OS **(E)** cells.

