## Supplementary Infomation for "E-cadherin represses anchorage-independent growth in sarcomas through both signaling and mechanical mechanisms"

^1^Center for Theoretical Biological Physics, Rice University, Houston, Texas, USA; ^2^Department of Medicine, Duke University Medical Center, Durham, North Carolina, USA; ^3^Department of Applied Physics, Rice University, Houston, Texas, USA; ^4^School of Medicine, Johns Hopkins University, Baltimore, Maryland, USA; ^5^Department of Pathology, Duke University Medical Center, Durham, North Carolina, USA; ^6^ Department of Bioengineering, Rice University, Houston, Texas, USA; ^7^Medical Scientist Training Program, Baylor College of Medicine, Houston, TX, USA; ^8^Department of Orthopaedics, Duke University Medical Center, Durham, North Carolina, USA; ^9^Flint Animal Cancer Center, Colorado State University, Fort Collins, Colorado, USA; ^10^Solid Tumor Program and ^11^Duke Prostate Center, Duke University Medical Center, Durham, North Carolina, USA; ^12^Department of Radiation Oncology, Duke University Medical Center, Durham, North Carolina, USA

#These authors contributed equally to this work.

**Supplementary Text**

**Supplemental Methods**

Model construction details

We developed a mathematical model that integrates our results with reported interactions, and models the regulations among E-cadherin, NRAGE, TBX2 and their effects on anoikis sensitivity. Our results show that E-cadherin overexpression downregulates TBX2 at mRNA level, suggesting an inhibitory link from E-cadherin to TBX2. Also, TBX2 has been shown to be a direct transcriptional repressor of E-cadherin (Wang et al., 2012). Further, E-cadherin can repress nuclear localization of NRAGE (neurotrophin receptor-interacting melanoma antigen) through ankyrin-G; nuclear localization can promote TBX2-mediated inhibition of anoikis (Kumar et al., 2011). The model formulation for the aforementioned interactions is given by:

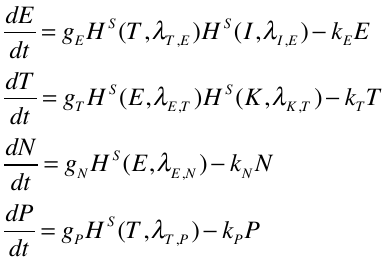

where E, T, N and P denote E-cadherin, TBX2, NRAGE, and p14ARF (surrogate measure of anoikis sensitivity in our model) levels respectively.
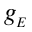
,
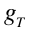
,
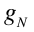
, and
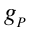
 are respective production rates for E-cadherin, TBX2, NRAGE, and p14ARF, and
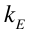
,
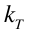
,
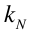
, and
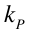
 are their respective degradation rates. Shifted Hill functions, defined as
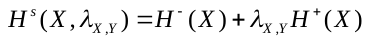
, denote the effect of X on Y, where
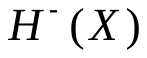
is the inhibitory Hill function,
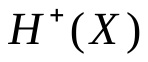
is the excitatory Hill function, and
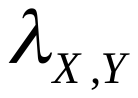
denote the equivalent of fold-change in production of Y due to X (Lu et al., 2013).
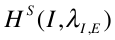
 captures the effect of overexpression of E-cadherin, and
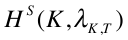
 represents TBX2 knockdown. Degradation rates (represented in per unit hour) for all proteins has been estimated based on typical half-life of proteins in humans, and the production rate (represented in molecules per hour) estimate is based on total number of protein molecules per cell reported for signaling molecules (~100,000) (Milo et al., 2010). Fold-changes in production have been estimated based on our results and existing literature: 0.4x for TBX2-mediated inhibition of E-cadherin (Wang et al., 2012), 0.4x for E-cadherin mediated inhibition of TBX2 (Fig 6A, B), and the inhibition of p14ARF by TBX2 and its accentuation by NRAGE (Kumar et al., 2011). All parameters are given in Table 1.

| Parameter | Value | Parameter | Value | Parameter | Value | Parameter | Value |
| --- | --- | --- | --- | --- | --- | --- | --- |
| 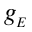 | 10000 | 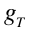 | 8000 | 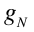 | 10000 | 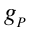 | 8000 |
| 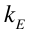 | 0.1 | 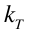 | 0.1 | 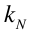 | 0.1 | 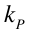 | 0.1 |
| 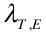 | 0.4 | 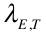 | 0.4 | 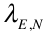 | 0.6 | 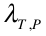 | 0.4 |
| 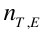 | 2 | 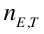 | 2 | 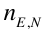 | 2 | 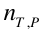 | 3 |
| 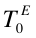 | 20000 | 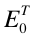 | 80000 | 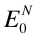 | 50000 | 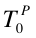 | 40000 |

**Table 1:** Parameters used to simulate effects of E-cadherin on anchorage-independent growth

The last two rows of parameters in Table 1 correspond to those included in shifted Hills function. A canonical inhibitory Hills function is defined as
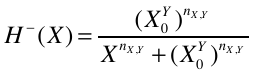
. To account for the case of NRAGE promoting the effect of TBX2 inhibiting p14ARF, the shifted Hills function for TBX2 inhibiting p14ARF is multiplied by (0.5+ (co/2)), where co =
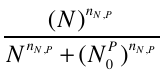
, such that the inhibition on p14ARF increases with increasing levels of NRAGE (with a usual saturation effect).

For external signals - overexpression of E-cadherin or knockdown of TBX2 – the value of signal is 5000 molecules, Hills coefficient =2, Hills threshold = 2000 molecules. Fold-change parameter (
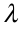
) was varied to mimic biological replicates, such as varying efficacy of siRNA-mediated knockdown. To calculate the fold-change in TBX2 or anoikis sensitivity (p14ARF) levels upon treatment with these external signals, we first calculated the steady state of the system without any external signal ([Ecad, TBX2, NRAGE, p14ARF] approximately equal to [45000, 70000, 85000, 40000] molecules), and then calculated the steady state of the system in the presence of external signal, for varying
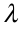
. Fold-changes observed in steady state values of p14ARF is reported here. The values of
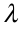
 considered pairwise for over-expression of E-cadherin and inhibition of TBX2 are [ (10, 0.4), (9, 0.45), (8, 0.5), (7, 0.55), (6, 0.6) ].

While considering the revised model (i.e. weaker inhibition of E-cadherin by TBX2), the following parameters are changed:
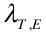
=
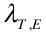
*2;
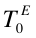
=
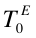
 *5. The new steady state of the system is calculated, and the fold-change observed in the steady state value relative to the new steady state is reported here. The same values of
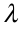
 as mentioned above are used.

**Supplemental Legends**

**Supplementary Figure 1. E-cadherin over-expression does not alter sarcoma cell growth in monolayer culture. A.** E-cadherin expression in RD cells was verified by immunofluorescence staining. **B.** E-cadherin expression had no effect on the in-monolayer growth rate of RD cells. **C.** E-cadherin expression in 143B cells was verified by immunofluorescence staining. **D.** E-cadherin expression had no effect on the in-monolayer growth rate of 143B cells. **E.** E-cadherin expression and membrane localization in U2OS cells was confirmed by immunofluorescence staining.

**Supplemental Figure 2. Ectopic E-cadherin expression in sarcoma cells does not alter EMT. A.** Ectopic expression of E-cadherin in RD cells has no influence on mesenchymal markers. **B.** E-cadherin expression has no effect on migration of RD cells. Images (left) were collected every two hours and quantified (right) using the IncuCyte Zoom system. **C.** E-cadherin expression does not change invasion in RD cells. **D.** Anchorage-independent growth of RD cells expressing E-cadherin was significantly inhibited. **E.**  Representative images of soft agar colony growth in RD cells with and without E-cadherin expression.

**Supplementary Figure 3. CREB knockdown indicates it is downstream of E-cadherin and TBX2. A.** CREB knockdown does not alter E-cadherin expression. **B.** CREB knockdown does not alter TBX2 expression.

**Supplementary Figure 4. E-cadherin induces E- to N-cadherin switching in sarcoma cells. A.** Immunofluorescence staining of E-cadherin+ clusters reveals a marked loss of N-cadherin,* indicates loss of N-cadherin expression. Conversely, E-cadherin- cells have strong N-cadherin staining at cell membranes. **B.** N-cadherin mRNA is not significantly different in E-cadherin-expressing cells. **C.** N-cadherin protein levels are unchanged in the presence of E-cadherin over-expression. **D.** N-cadherin (CDH2) knockdown was confirmed by immunofluorescence staining using two independent siRNAs. **E-G.** N-cadherin (CDH2) knockdown had no effect on anoikis resistance (**E**) or p-CREB protein levels (**F,G**).

**Supplementary Figure 5. Revised E-cad/TBX2 signaling model**. **A.** Weakened inhibition of E-cadherin by TBX2 denoted by dotted line. **B.** Revised mathematical model predicts relatively weak effect of the combinatorial treatment (TBX2-KD + Ecad-OE) as compared to only one treatment (Ecad-OE or TBX2-KD) as compared to that seen in Fig 6B.

**Supplementary Figure 6. A mechanical model relates cell-cell adhesion to spheroid size. A.** As adhesion increases, spheroid size becomes smaller. **B.** The model predicts that cell-cell adhesion is positively correlated with the average number of neighbors for each cell in a spheroid.
